# Goats who stare at wolves - identifying natural response stimuli for an affect-driven attention bias test in small ruminants

**DOI:** 10.64898/2025.12.03.692016

**Authors:** Jana Deutsch, Steve Lebing, Anja Eggert, Christian Nawroth

## Abstract

Reliable and non-invasive assessment of affective states is crucial in animal welfare research. Linking affective states to cognitive processes, such as cognitive biases, provides a promising approach. An affect-driven attention bias - the tendency to focus more on threatening than on neutral or positive stimuli when in a negative affective state - has been observed in humans and non-human animal species. However, implementation is often hampered because ecologically relevant test setups need to be adapted to species-specific differences in the perception and processing of information from different modalities. As an initial step towards developing and validating an affect-driven attention bias test for goats, we aimed to identify visual stimuli perceived as potentially threatening using a looking-time paradigm. In a within-subject design, 30 adult female dwarf goats (*Capra hircus*) were presented with photographs of 12 animal species (six natural predators and six non-predators) from three taxa (mammals, reptiles, birds). Each image was shown for 10 seconds on one of two video screens, while the opposite screen remained white. Each goat completed four sessions (one per day) of six trials each: two sessions showing full-body and two sessions showing face/head-only images, resulting in 24 trials per subject. Analysis of relative looking duration revealed a significant interaction between animal taxon and predator category. Goats looked longer at predatory than non-predatory reptiles, while the opposite pattern was found for mammals. No difference was observed for birds. The presented body part (full body vs. face/head only) did not influence looking behaviour. These results suggest that photographs of predatory reptiles, particularly snakes, might be perceived as potentially threatening by goats, indicating their suitability as negative stimuli in future affect-driven attention bias tests. Further validation is needed to confirm their negative valence.

## 1. Introduction

Since the 1960s, animal welfare researchers have been addressing the question of whether farmed animals are doing well (e.g. Harrison, 1964), and the issue has long reached the centre of society (e.g. Alonso et al., 2020; Bennett, 1995; Miele et al., 2011). While the precise definition of “animal welfare” has been strongly debated since then (for a detailed elaboration on different welfare concepts, see Fraser, 2008), researchers nowadays mostly agree on the assumption that it requires more than solely the fulfilment of basic needs regarding health and housing conditions (Mellor, 2016; Rault et al., 2025). For example, most current frameworks consider the affective states of animals as an important component of animal welfare (Fraser, 2008; Mellor, 2016; Rault et al., 2025). These affective states can occur as short-term emotions or long-term moods (Mendl et al., 2010). While emotions are multi-component responses to specific events that are relevant from a survival perspective, moods are defined as “free-floating valenced states” that are not directly tied to a specific event but arise due to the cumulative experience of emotions (Mendl & Paul, 2020). According to the “core affect” model, affective states are generally classified and quantified along at least two dimensional scales, one that identifies valence (i.e. whether a state is hedonically positive or negative in terms of pleasant or unpleasant) and one that characterises the arousal level (i.e. one that refers to physiological activation) of the state (Mendl & Paul, 2020; Paul et al., 2005; Russell, 2003). In humans, affective states are often assessed via verbal self-report, making the transfer to non-human subjects, such as farm animals, challenging (Mendl et al., 2022).

Affective states in non-human animals can, for example, be assessed using physiological measures such as heart rate variability (e.g. Krause et al., 2017), or behavioural parameters such as vocal (e.g. Leliveld et al., 2016) and facial expressions (e.g. Finlayson et al., 2016). Nevertheless, many commonly used parameters of affective states might be stronger associated with arousal than valence (Paul et al., 2005). An animal’s affective state, particularly its valence, can also be evaluated by linking it to cognitive processes. One example of how affective states are intertwined with how information is processed are so-called cognitive biases (Harding et al., 2004; Mendl et al., 2009). Studying cognitive processes related to decision-making or attention in non-human animals, thus, offers a promising approach to draw conclusions about their underlying affective states. One established method to do this is the judgment bias test (JBT). In this paradigm, animals are trained that one cue predicts a positive event, while another predicts a less positive or even negative event. After initial training, subjects are presented with intermediate (and therefore ambiguous) stimuli and their reactions to these cues are analysed as a proxy for the underlying affective state. A subject in a more positive affective state is assumed to be more optimistic in these ambiguous situations compared to a subject in a negative affective state (Harding et al., 2004). The JBT is a very popular cognitive bias test in non-human animals (e.g., see Burman et al., 2008 for JBT in rats (*Rattus norvegicus*); Düpjan et al., 2017 for JBT in pigs (*Sus scrofa*); Neave et al., 2013 for JBT in cattle (*Bos taurus*)) and has often proven to be a suitable tool for assessing affective states (Ede & Parsons, 2023; Lagisz et al., 2020; Paul et al., 2022). However, its implementation comes with certain disadvantages, such as the need for extensive animal training, which is both time- and cost-intensive. Another weakness of the JBT is the difficulty of repeatedly testing the same subjects with the same experimental setup, as subjects will rapidly learn that the intermediate cues are often not rewarded, which may affect their response to them (e.g., see Doyle et al., 2010), raising doubts about whether their reactions accurately reflect the underlying affective state.

An alternative approach to assessing cognitive biases focuses on attention rather than judgment. Attention can be biased in terms of how attentional resources are allocated towards one stimulus compared to others (Crump et al., 2018). In the human literature, attention biases modulated by external factors of a stimulus (e.g., see Horstmann & Herwig, 2016 for an attention bias to novelty) as well as attention biases towards or away from emotional information influenced by the observer’s affective state (so-called “affect-driven attention biases”, hereafter ADAB) are reported. For instance, an ADAB can be observed in clinically anxious participants, who show increased attention towards threat-related stimuli in a probe detection paradigm compared to non-anxious control participants (MacLeod et al., 1986). As attention allocation can, for example, be assessed via looking duration at specific stimuli (Crump et al., 2018; see Kappel et al., 2025 for a detailed discussion of vision-based ADAB test designs), little task-related training is needed, making ADAB tests time-efficient and therefore potentially feasible for on-farm application. These features make ADAB tests, originally derived from human clinical studies, suitable candidates for future use in assessing affective states in non-human animals while overcoming some of the disadvantages of the JBT.

ADAB tests have already been used to assess affective states in a set of non-human animals. Bethell et al. (2012), for example, presented captive rhesus macaques (*Macaca mulatta*) with pictures of conspecific faces showing an aggressive expression following a veterinary health check (stress condition) and during a period of environmental enrichment (neutral or potentially positive condition), and recorded their looking behaviour towards the presented stimuli. While the aggressive faces were actively avoided in the stress condition, subjects sustained higher attention to this stimulus category during the enrichment period, indicating that a shift in emotional state can affect social attention in non-human primates (Bethell et al., 2012). Campbell et al. (2019) showed in an ADAB test for chickens *(Gallus domesticus)* that laying hens that never ranged were more vigilant and slower to feed after being confronted with a conspecific alarm call (simulating a threat) compared to hens that ranged daily, indicating higher anxiety in indoor-preferring hens. In pharmacologically treated high (m-chlorophenylpiperazine) anxiety sheep (*Ovis aries*), increased attention to a live dog (presented for 10 seconds as a threat stimulus) and increased vigilance were shown compared to low (diazepam) anxiety sheep (Lee et al., 2016). This study also assessed the latency to feed after being confronted with the threat stimulus, therefore using a feed reward as a positive stimulus. However, when comparing pharmacologically induced anxious (m-chlorophenylpiperazine), calm (diazepam), happy (morphine) and control (saline) sheep, no significant differences were found between treatment groups for the duration of vigilance or looking behaviours to the stimuli presented in the ADAB test, even though anxious subjects tended to be more vigilant than control animals and had a longer latency to become non-vigilant (Monk et al., 2019a). In the latter test, a life-size photograph of a conspecific was used as a positive stimulus, while a live dog presented for 3 seconds was presented as a threat stimulus, indicating that the replicability of results might be strongly influenced by stimulus choice.

When conducting ADAB studies, researchers must carefully consider the choice of stimuli, their valence, and how they are perceived by the animal species under investigation. While in some ADAB studies auditory stimuli like conspecific alarm calls were used as negative stimuli (e.g., see Brilot & Bateson, 2012 (starlings (*Sturnus vulgaris*)); Campbell et al., 2019 (chickens)), in others subjects were confronted with live stimuli like an unfamiliar human (Cussen & Mench, 2014 (Orange-winged amazon (*Amazona amazonica*))) or a dog (Lee et al., 2016; Monk et al., 2018a, 2018b, 2019a, 2019b (sheep); Lee et al., 2018 (cattle)). Alternatively, video stimuli of conspecifics showing agonistic behaviour (e.g., Vögeli et al., 2015 (sheep)) or images of naturally aversive eyespots (Brilot et al., 2009 (starlings)), a veterinarian (Allritz et al., 2016 (chimpanzees (*Pan troglodytes*))) or aggressive conspecific faces (Bethell et al., 2012 (rhesus macaques)) were presented as threat stimuli. Stimulus characteristics (2D vs. 3D, still images vs. video recordings, etc.) and modalities (visual, acoustic, etc.) can influence how subjects respond to them. Thus, stimulus validation, likewise in terms of their respective valence, is a key step when developing ADAB tests.

To identify images that could subsequently be presented as negative stimuli against neutral or positive ones in a potential future ADAB test, we analysed behavioural responses to ecologically relevant visual stimuli using a looking time paradigm (Deutsch et al., 2024). We decided to use visual stimuli for ADAB testing as they might be the best option with regard to feasibility and scalability for later on-farm application. In primates, visual detection of pictures of snakes is enhanced, indicating an innate attentiveness to this natural predator class to avoid predation risk (e.g., see Shibasaki & Kawai, 2009). Cattle increased vigilance and decreased foraging rates when presented with three-dimensional visual and olfactory stimuli of wolves (Kluever et al., 2009). Raptor-shaped images presented on an overhead video screen have been shown to elicit alarm calls and non-vocal anti-predator behaviour in chickens (Evans et al., 1993). As we aimed at identifying visual stimuli that goats naturally perceive as threatening and that would not require additional negative conditioning, we decided to present photographs of animal species that are either natural predators of goats (in the case of canids), goat kids (in the case of raptors) or species that might otherwise be perceived as dangerous to them (in the case of venomous snakes). The species we presented naturally occur (at least on the taxonomic family level) in the Fertile Crescent, a region in the Middle East in which early human civilisations domesticated goats around 10,000 years ago (Luikart et al., 2001). As a control, we also presented non-predatory species, likewise occurring in the Fertile Crescent. We presented dwarf goats with the images in a looking time task, and analysed the looking behaviour towards the different stimuli spanning threat categories (predators or non-predators), animal taxa (mammals, birds and reptiles), and body parts (full body or head/face only depicted). We hypothesised that goats would show different behavioural responses to two-dimensional images of predators compared to non-predators, irrespective of the taxon of the presented stimulus species. We therefore predicted that the goats in our study would look longer and from an increased distance at predator compared to non-predator stimuli, indicating an increased attention to this stimulus type while in a state of alertness and readiness to escape.

## 2. Animals, Materials and Methods

### 2.1 Ethical note

The study was waived by the State Agency for Agriculture, Food Safety and Fisheries of Mecklenburg-Vorpommern (Process #7221.3-18196_24-2) as it was not considered an animal experiment in terms of sect. 7, para. 2 Animal Welfare Act. Animal care and all experimental procedures were in accordance with the ASAB/ABS guidelines for the use of animals in research (ASAB Ethical Committee/ABS Animal Care Committee, 2024). All measurements were non-invasive, and the experiment did not last longer than ten minutes per day for each goat. The test would have been stopped if the goats had shown signs of an increased stress level.

### 2.2 Subjects and Housing

Thirty non-lactating female Nigerian dwarf goats reared at the Research Institute for Farm Animal Biology (FBN) in Dummerstorf participated in the experiment. The subjects were between one and four years old at the start of data collection (Groups A and B: one year old; groups C and D: two years old; group E: between three and four years old; see S1 for detailed information on test subjects) and were kept in five groups with six animals each. All animals were experimentally pre-experienced, although the degree of experience varied (see S1). Each group was housed in an approximately 15 m^2^ (4.8 m x 3.1 m) pen consisting of a deep-bedded straw area (3.1 m x 3.1 m) and a 0.5 m elevated feeding area (3.1 m x 1.5 m). Each pen was equipped with a hay rack, a round feeder, an automatic drinker, a licking stone, and a wooden podium for climbing. Hay and food concentrate were provided twice a day at 7 am and 1 pm, while water was offered ad libitum. Subjects were not food-restricted during the experiments.

### 2.3 Experimental arena and apparatus

The experimental arena was located next to the home pens. It consisted of three adjoining rooms with 2.1 m high wooden walls connected by doors (see S2). Data collection took place in a testing area (4.5 m x 2 m) divided into two parts (2.25 m x 2 m) by a fence that facilitated the separation of single subjects from the rest of the group. The experimental apparatus was inserted into the wall between the testing area and the experimenter booth (2 m x 1.5 m), which was located behind the apparatus and where an experimenter (E1) was positioned during all sessions. The subject in the testing area had no visual contact with E1. Since testing was conducted individually, the rest of the animal group remained with the other experimenter (E2) in an adjacent waiting area (6 m x 2.2 m).

The experimental apparatus (Fig. 1; previously described in Deutsch et al., 2024) was inserted into the wall between the testing area and the experimenter booth at a height of 36 cm above the floor and consisted of two video screens (0.55 m x 0.33 m) mounted on the rear wall of the apparatus. The video screens were positioned laterally so that they were angular (around 45°) to a subject standing in front of the apparatus. Regarding height, subjects standing in front of the apparatus were considered to be looking approximately at the centre of the screens. A digital camera (AXIS M1124, Axis Communications, Lund, Sweden) was installed between the upper edge of the wall separating the two video screens and the ceiling of the apparatus. This provided an angled view of the subject inside the apparatus and the space in proximity to the apparatus in the testing area. Videos were recorded with a 30 FPS rate. A food bowl, connected to the experimenter booth by a tube, was inserted into the bottom of the apparatus. This allowed E1 to deliver food items without being in visual contact with the test subject. As an alteration to the experimental apparatus described in Deutsch et al. (2024), a wooden board with a round cutout at the height of the animal’s neck and a plexiglass pane was attached to the apparatus to prevent the animals from climbing into it.

**Fig. 1.**
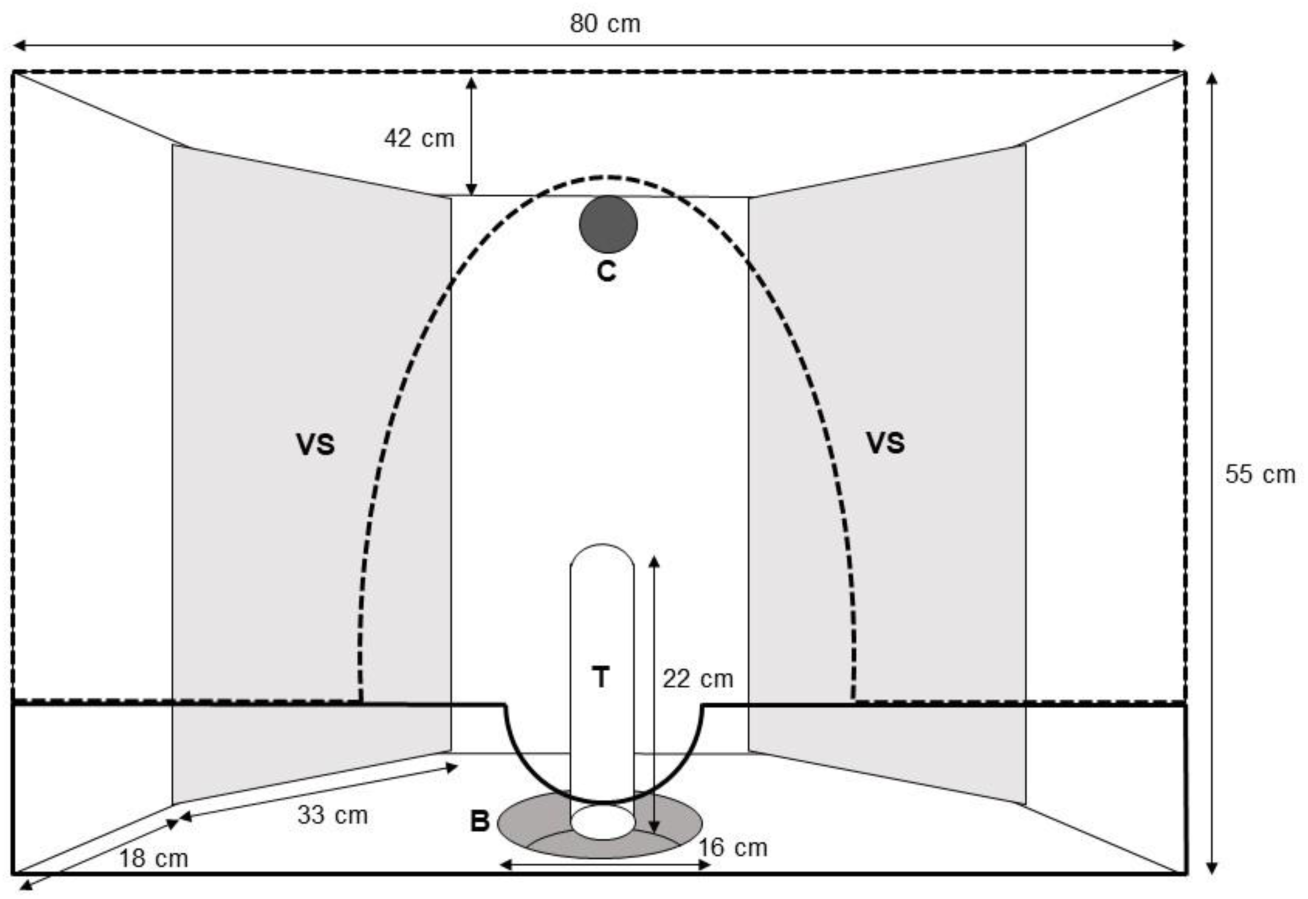
Experimental apparatus with video screens (VS), camera (C), food bowl (B), tube (T), wooden board with a round cutout (solid line) and a plexiglass pane (dashed line)

### 2.4 Habituation

Habituation was conducted following the habituation protocol described in detail in Deutsch et al. (2024). The duration of the single habituation steps was adjusted for each group since they differed in their experimental pre-experience, including their familiarity with human handlers and the experimental apparatus (groups C-E had previous experience with this experimental setup, while groups A and B were naive; see S3 for further details). Briefly summarised, the habituation steps were the following: Habituation to human handlers (here the experimenters E1 and E2) in the home pen, habituation to the experimental arena as a whole group, habituation to the experimental apparatus as pairs and habituation to the experimental apparatus as individuals (see S4 for the objectives of the single habituation steps). Habituation for groups A and B took place from March to May 2024, for groups C and D in June 2024 and for group E in July 2024. At the end of each habituation period, all animals remained calm in the testing area and fed from the food bowl and, therefore, all thirty subjects proceeded to the data collection during which one animal had to be excluded at a later stage as it began to show indicators of high stress (e.g. increased vocalisations, restless wandering, and rejection of feed intake).

### 2.5 Experimental procedure

#### 2.5.1 Stimuli and stimulus presentation

Photographs of twelve animal species (see S5) were used as stimuli. Photographs were obtained from a royalty-free online platform (https://pixabay.com/de/) and then modified to remove the background (GIMP 2.10.38, GNU Image Manipulation Program, Copyright © 1995-2024, Spencer Kimball, Peter Mattis and the GIMP development team). The presented species belonged to three animal taxa (mammals, birds and reptiles) with four species each. Half of the stimuli (wolf, jackal, red kite, eagle, viper and cobra) were categorised as potential threat stimuli as they represented either natural predators of adult goats (in the case of the wolf and the jackal), natural predators of goat kids (in the case of the red kite and the eagle) or animals that might be harmful to goats (in the case of the viper and cobra). The other half of the stimuli (donkey, antelope, duck, pigeon, chameleon and gecko) were categorised as potential neutral stimuli and were matched to the predatory species regarding their taxon (see Fig. 2 for an example). Each species was presented in a test trial either as a full-body photograph or a photograph showing the face/head only. The brightness and size of each stimulus image were assessed (ImageJ 1.53m, Wayne Rasband and contributors, National Institute of Health, USA, https://imagej.net, Java 1.8.0-internal (32-bit)). Stimuli did not substantially differ concerning brightness (predatory mammals: 117 ± 17.0 (mean ± SD), non-predatory mammals: 130 ± 17.3, predatory birds: 102 ± 21.6, non-predatory birds: 119 ± 20.0, predatory reptiles: 134 ± 14.8, non-predatory reptiles: 128 ± 9.3) and size (predatory mammals: 151855 ± 12903 px (mean ± SD), non-predatory mammals: 188120 ± 39515 px, predatory birds: 169226 ± 35616 px, non-predatory birds: 140940 ± 27752 px, predatory reptiles: 171908 ± 35175 px, non-predatory reptiles: 168483 ± 44918 px). The stimuli were presented in colour in front of a white background. Stimuli were presented either on the left or the right screen while the other screen remained white. Each test session consisted of a stimulus set containing six stimuli, either showing only full-body photographs or face/head-only photographs. Each of these sets contained photographs of one predatory and one non-predatory species of each taxon, with each stimulus being presented only once (see S6 for detailed information on the stimulus arrangement in the sets). The stimuli were presented on the video screens in a pseudorandomised and counterbalanced order.

**Fig. 2.**
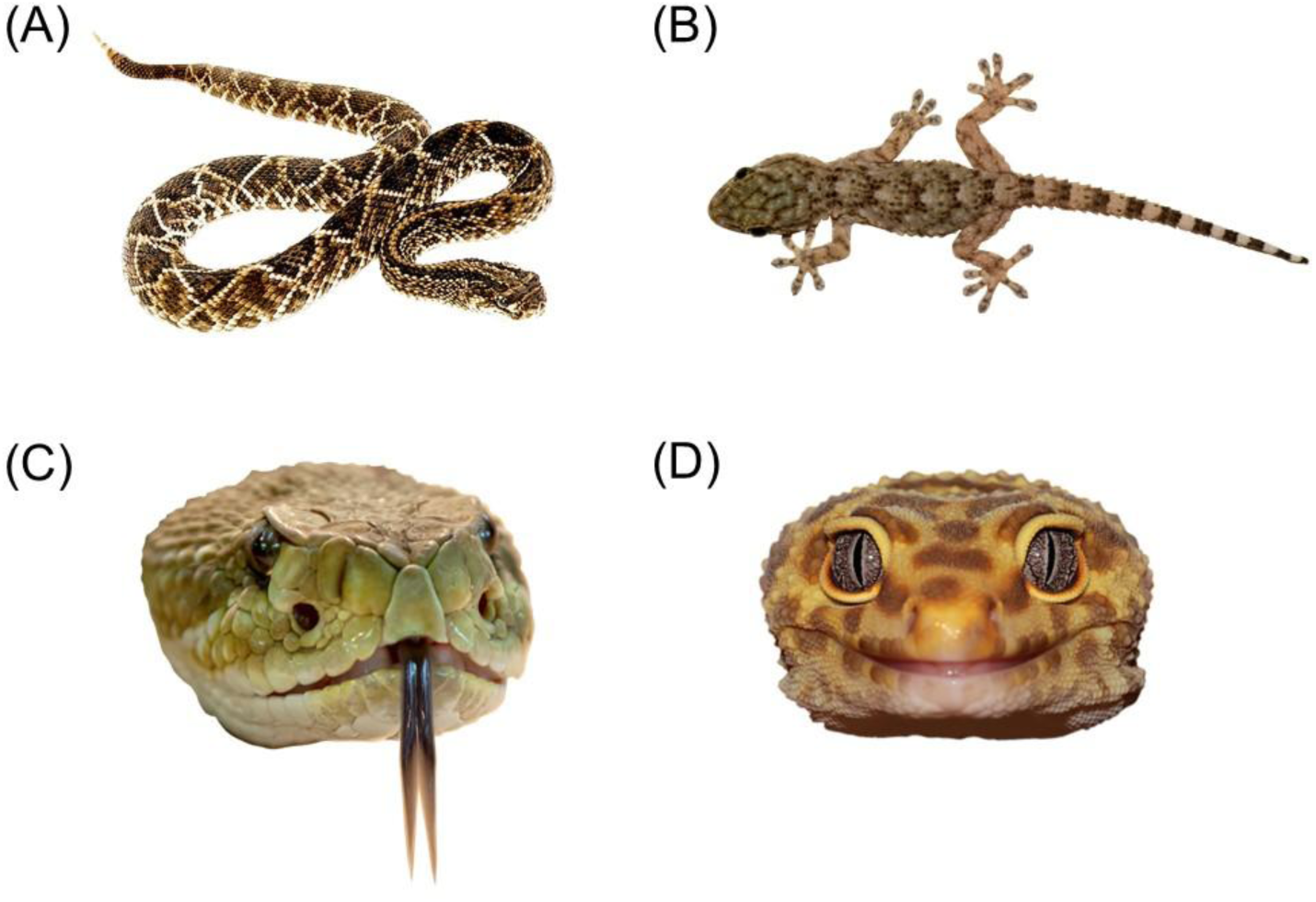
Example of the photographs used as stimuli (A) full-body photograph of viper, (B) full-body photograph of gecko, (C) face photograph of viper and (D) face photograph of gecko

#### 2.5.2 Data collection

Data collection took place from May 2024 to July 2024: groups A and B were tested in May, groups C and D in June, and group E in July. All procedures followed the testing protocol described in Deutsch et al. (2024). Testing started between 8:00 and 09:00 a.m. each day. Each subject completed four sessions of six trials each (two sessions with full-body stimuli and two with face stimuli), with one session conducted per day (see S7 for subject set assignment).

A session started when the subject was separated from the rest of the group and stood in front of the experimental apparatus. Prior to the stimulus presentation, one to two motivational trials were conducted, during which a food item was inserted through the tube into the food bowl without any stimulus being presented for 10 seconds. Immediately before each stimulus presentation, a food item was again inserted into the food bowl, followed by a 10-second stimulus presentation. Motivational and test trials alternated until all six stimuli of a set had been presented. The number of motivational trials varied depending on the behaviour of the subject and could be increased, e.g. if the animal was restless at the beginning of the session. Data from the subject that was excluded after the second test session remained in the dataset.

### 2.6 Data scoring and analysis

#### 2.6.1 Video coding

The behaviour of the individual goats was scored using Boris (Friard & Gamba, 2016, Version 8.27.1), an event logging software for video coding and live observations. Video coding was performed in frame-by-frame mode for the ten seconds of stimulus presentation per trial. Two main parameters were scored: (1) the duration for which the subject’s head was visible inside the experimental apparatus or within a defined area near it (see Fig. 3), and (2) the duration spent looking at each video screen from inside and outside the apparatus. A subject’s head was considered ‘visible’ when two conditions were met: both forelegs (i.e., both complete claws) had crossed a fictitious line that defined the scoring area (see Fig. 3; the area was defined as the looking direction of the animals could be accurately determined up to this distance) and the observer could determine whether the subject was looking at a screen or not. To determine the subject’s looking direction, a second fictitious line that extends from the middle of the snout (orthogonal to the line connecting both eyes) was drawn (Fig. 3). The alignment of this line with the subject’s binocular focus served as an indicator of attention directed towards a particular screen. Looking duration was only scored when the head was scored as ‘visible’. Besides the total looking duration at a specific screen, the distance from which the stimulus was observed (from inside or outside the apparatus) was also scored. This distinction was made on whether the tip of the snout crossed the wooden board in the apparatus while the subject was looking at a screen. Looks were scored as ‘inside’ when the snout crossed the board, and as ‘outside’ in all other cases. Inter-observer reliability for looking duration was assessed and found to be very high (72 out of 719 trials (10 %) of the videos were coded independently by two observers, yielding a Spearman’s correlation coefficient of ρ = 0.97; p < 0.001).

**Fig. 3.**
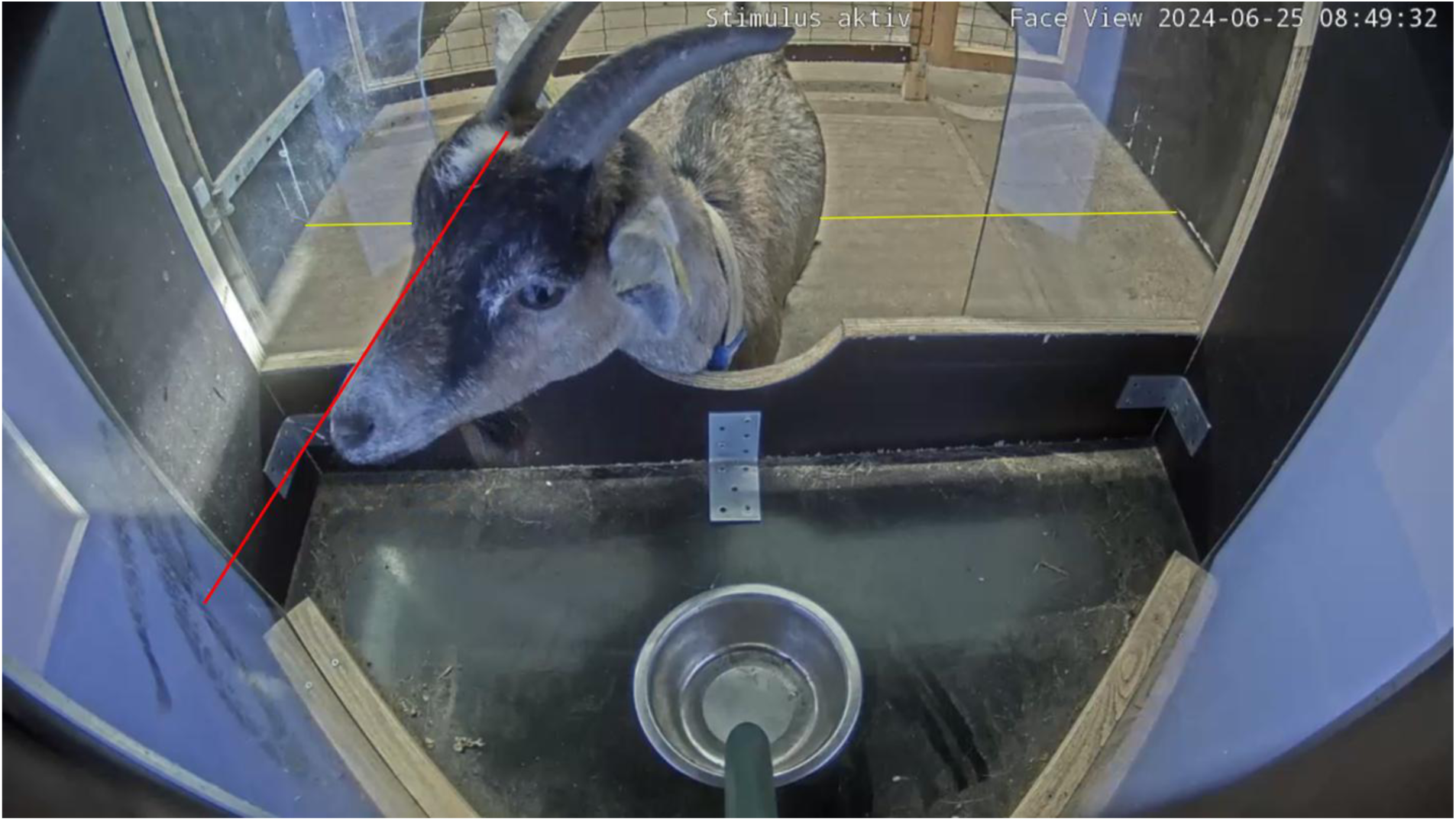
Image of the camera perspective used for video coding showing the scoring area (area in front of the yellow line). The head was scored as “visible” when both complete claws of the forelegs crossed the line, and the observer could determine whether the subject was looking at a screen or not. Fictitious line extending from the middle of the snout (red) that was used to determine the subject’s looking direction. In this example image, the animal’s head would be scored as “visible” and its looking direction as “right” (from the perspective of the subject) and “inside the apparatus” as the tip of the snout had crossed the wooden board in the apparatus

#### 2.6.2 Statistical analysis

All statistical analyses were conducted in R (Version 4.4.2; R Core Team, 2024).

##### 2.6.2.1 Analysis of relative looking duration

To examine factors influencing the duration goats spent looking at the video screen presenting a stimulus (S+), we fitted a Bayesian linear mixed-effects model (BLMM) using the R package “blme” (Chung et al., 2013). The response variable, “Relative looking duration at S+” (RLD) calculated as:

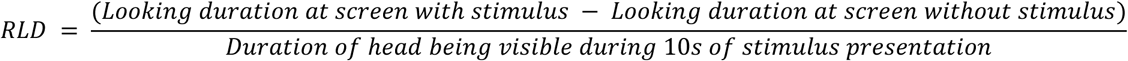

Trials with a looking duration of zero at the stimulus screen (n = 27) were excluded, as such cases likely indicated distraction. The model included “Category” (two levels: predator, non-predator), “Taxon” (three levels: mammal, bird, reptile) and “Body part” (two levels: full-body, head/face only) as fixed effects, as well as all two-way interactions among these factors. Random effects were specified as “Session” (1-4) nested within “Subject” (goat identity), which in turn was nested within “Group” (A-E). Model assumptions were checked using graphical residual inspection and simulation-based tests with the “DHARMa” package (Hartig et al., 2024). Global model significance was assessed using parametric bootstrapping (1,000 simulations) using the “pbkrtest” package (Halekoh & Højsgaard, 2014). When the global model was significant, individual predictors (including their interactions) were tested by comparing the full model with reduced models omitting the respective predictor. Parametric bootstrap tests were preferred over raw likelihood ratio tests (LRTs) as they do not rely on large-sample asymptotic approximations and correctly account for complex random-effect structures (Halekoh & Højsgaard, 2014).

##### 2.6.2.2 Analysis of looking distance

To analyse which factors influenced the distance from which goats observed the stimuli (i.e. from inside vs. outside the apparatus), we fitted a generalised linear mixed-effects model (GLMM) with a beta error distribution and logit link using the “glmmTMB” package (Brooks et al., 2017). The response variable, “Relative looking duration at S+ from outside” (RLD_Out), was calculated as:

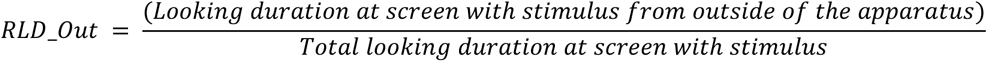

As in the BLMM, trials with zero looking duration were excluded. The GLMM included the same fixed and random effects as the BLMM. Model assumptions were evaluated using simulation-based residual diagnostics (“DHARMa”; Hartig et al., 2024). Overdispersion and underdispersion did not affect model results. Model significance was again assessed through parametric bootstrapping (1,000 samples), using likelihood ratio tests (LRTs) between full and reduced models. For each bootstrap iteration, new response data were simulated from the reduced model, both models were refitted, and an empirical distribution of LRT values was obtained. The p-value was computed as the proportion of simulated LRT values greater than or equal to the observed LRT value. This procedure paralleled that used for the BLMM, but was adapted here for the GLMM framework.

## 3. Results

The relative looking duration (RLD) was significantly influenced by the taxon of the presented stimulus species (p = 0.001). In addition, a significant interaction was found between “Category” (predator vs. non-predator) and “Taxon” (p = 0.016). No significant main effects of “Body part” (p = 0.718) or “Category” (p = 0.706), nor interactions involving “Body part” (Category x Body part, p = 0.297; Taxon x Body part, p = 0.285), were found.

Visual inspection of the model estimates (Tab. 1) and visual representation of the data (Fig. 4) indicated that goats looked longer at predatory reptiles compared with non-predatory reptiles, whereas the opposite pattern was observed for mammals, with longer looking durations towards the non-predatory species. No meaningful difference in looking duration at predatory and non-predatory birds was observed. However, given the overlapping confidence intervals among stimulus classes, these tendecies should be interpreted cautiously.

**Tab. 1.** Table containing the model estimate, lower confidence interval and upper confidence interval for each presented stimulus class

| Category | Taxon | Body part | Estimate | Lower CI | Upper CI |
| --- | --- | --- | --- | --- | --- |
| Non-predator | Bird | Face | 0.209 | 0.099 | 0.324 |
| Predator | Bird | Face | 0.193 | 0.084 | 0.306 |
| Non-predator | Bird | Body | 0.206 | 0.091 | 0.322 |
| Predator | Bird | Body | 0.218 | 0.102 | 0.335 |
| Non-predator | Mammal | Face | 0.203 | 0.089 | 0.32 |
| Predator | Mammal | Face | 0.136 | 0.031 | 0.249 |
| Non-predator | Mammal | Body | 0.185 | 0.075 | 0.297 |
| Predator | Mammal | Body | 0.147 | 0.04 | 0.263 |
| Non-predator | Reptile | Face | 0.248 | 0.139 | 0.364 |
| Predator | Reptile | Face | 0.276 | 0.167 | 0.387 |
| Non-predator | Reptile | Body | 0.196 | 0.084 | 0.308 |
| Predator | Reptile | Body | 0.253 | 0.136 | 0.364 |

**Fig. 4.**
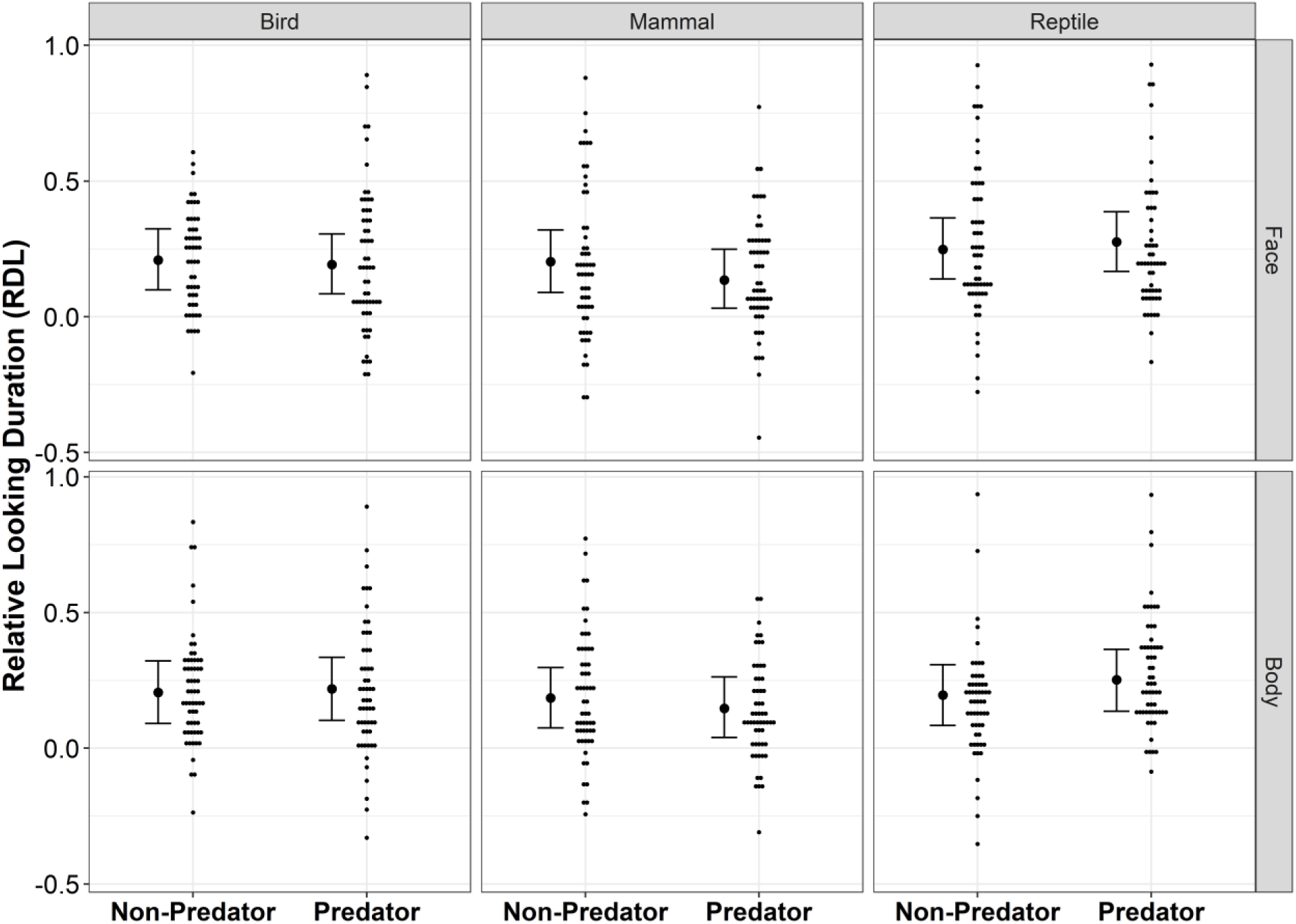
Relative Looking Duration of the subjects across taxon, category and body part of the presented stimulus species. Small black dots represent raw data points. Black dots plus lines and whiskers are the corresponding model estimates for each condition, and the 95% confidence intervals of the maximum model, respectively The relative looking duration at S+ from outside (RLD_Out) was not significantly influenced by Taxon (p = 0.248), Body part (p = 0.285), or Category (p = 0.217). Moreover, no significant interaction effects were detected (all p > 0.05; for detailed information on model estimates, and lower and upper CIs for each presented stimulus class, see S8).

## 4. Discussion

Attention bias testing has the potential to be a powerful approach for assessing affective states in non-human animals. However, test protocols must be adapted to each species. In this study, we tested whether photographs of natural predators of goats could serve as negative stimuli in future affect-driven attention bias (ADAB) tests that would not require additional negative conditioning. Selecting species-specific stimuli that actually elicit the expected perceived valence is a crucial step in developing valid ADAB tests (Kappel et al., 2025). Dwarf goats were presented with images in a looking time task, and the looking behaviour towards different stimuli spanning threat categories (predators or non-predators), animal taxa (mammals, birds and reptiles), and body parts (full body or head/face only depicted) was analysed. We found that looking behaviour towards predators and non-predators depended on the taxon of the presented stimulus species. Our results therefore add relevant information on stimulus choice when adapting ADAB tests to small ruminants, enabling species-specific adaptation to its design.

Visual attention of the subjects to predator and non-predator stimuli was influenced by the presented animal taxon. Therefore, we need to reject our hypothesis that goats would show different behavioural responses to two-dimensional images of predators compared to non-predators, regardless of the taxon of the presented stimulus species. Furthermore, our prediction that the goats would look longer at predator stimuli compared to non-predator stimuli, indicating increased attention towards threat-related cues, was not supported, as this was only evident for the reptile stimuli. Goats spent more time looking at predatory reptiles than at non-predatory reptiles. A possible explanation for directing more attention to the images of vipers and cobras might be an innate high attentiveness to snakes, which has been observed in humans (Bertels et al., 2020; Kawai, 2019) and non-human primates (Shibasaki & Kawai, 2009). Although extensive research has focused on innate fear responses to visual snake cues in humans and non-human primates, evidence for other mammalian species is still scarce. While there is no research on this topic in ungulates, some studies on rodents have examined fear-like responses elicited by snake-related stimuli (e.g. Guimarães-Costa et al., 2007; Watanabe et al., 2022). For example, Watanabe et al. (2022) found that degus (*Octodon degus*) exhibited visually induced antipredator behaviour in response to photographs of snakes, despite having been born and raised in a laboratory environment without prior exposure to snakes. In contrast, mice (*Mus musculus*) did not show snake aversion in the same study. Our results may therefore provide the first indication of a similar behavioural pattern in a farm animal species. However, further research is required to confirm this interpretation.

Unexpectedly, goats did not pay more visual attention to the canines presented as predatory mammals compared to the ungulates presented as non-predatory mammals. This might suggest that goats did not perceive the wolf and jackal stimuli as potential threats. Some animals (groups C–E) had been allowed to roam freely on a pasture prior to data collection and may therefore have encountered domestic dogs in the vicinity of the pasture. This might have led to a general mental categorisation of canines, regardless of species, as “non-threatening”. Alternatively, goats may have shown a visual preference for ungulates over canines because of morphological similarity. Among the non-predatory species presented, antelopes and donkeys most closely resemble goats and may subsequently have attracted more attention. If the ungulate stimuli were perceived as “goat-like”, this could have produced a stress buffering effect, as the animals were isolated from the rest of their social group during testing. Similar stress buffering effects of conspecific photographs have been demonstrated in isolated sheep (Da Costa et al., 2004).

The mode and duration of stimulus presentation likely also plays a critical role in reliable ADAB testing. Live dogs have repeatedly been used as threat stimuli in ADAB tests with sheep (Lee et al., 2016; Monk et al., 2018a, 2018b, 2019a, 2019b) and also in a study on the cardiovascular and behavioural effects to dogs in pregnant and lactating goats (Olsson & Hydbring-Sandberg, 2011). Even though dogs generally seem to be perceived as a threat by small ruminants (e.g. Lee et al., 2016; Monk et al., 2018b; but see Raoult & Gygax, 2018 for contrasting findings with video stimuli of dogs in sheep), reactions to the canines were not always consistent. For example, anxious sheep showed an attention bias away from a briefly (3 seconds) presented dog towards photographs of conspecifics (Monk et al., 2018a). In contrast, anxious sheep showed an attention bias to the dog (here presented for 10 seconds) away from a food bowl in another study (Lee et al., 2016). It is important to notice that in these studies, attention biases were assessed as the behavioural reactions after removal of the threat stimulus, while our study measured visual attention during exposure to the stimuli. Therefore, it remains unclear whether canines consistently elicit fear responses in small ruminants. The mode and duration of stimulus presentation, as well as the presence of competing positive cues, are very likely to influence behavioural reactions in attention bias tests.

Moreover, in our study, goats showed no differential looking behaviour towards predatory and non-predatory birds. Several factors might explain this finding. First, birds of prey are mainly predators of goat kids, and as our test subjects were adults, they may have lost any potential fear reaction towards raptors after their juvenile stage. Additionally, the presentation method may have reduced ecological relevance. From a natural perspective, raptors would typically be perceived by goat kids as overhead silhouettes during an attack. To reflect this, we presented images of birds in flight as seen from below. However, since the stimuli were displayed on video screens positioned in front of the subjects, their actual ecological relevance might have been limited. Furthermore, facial photographs of birds are unlikely to represent meaningful stimuli for prey animals, as a raptor’s face is rarely seen from the ground. An overhead presentation of bird photographs might therefore increase their ecological validity (e.g., see Evans et al., 1993).

We also analysed the distance from which stimuli were observed, assuming that it would be a suitable additional indicator for the perceived valence of the stimuli. We expected that the goats would maintain a greater distance from the predator stimuli compared to the non-predators, as it is reasonable for prey animals to keep a safe distance when encountered with a threatening animal. Contrary to our expectations, the type of stimuli did not affect the distance from which they were observed. This result likely reflects similar limitations in stimulus salience and presentation methods discussed above.

Stimulus presentation itself represented one of the main challenges in this study. To maintain ecological validity as much as possible, we used colour images. However, as the selected stimulus species differed slightly in their coat, skin, or plumage colouration, we cannot exclude that looking behaviour was influenced by image properties such as colour or contrast. Furthermore, the goats might not have perceived the two-dimensional images on a video screen as representations of real animals. Contrary to this, Langbein et al. (2023) provided some evidence that goats may associate two-dimensional photographs of conspecifics with their real-life counterparts. Another approach could be to use video clips instead of still images, which might provide more ecologically relevant stimuli and elicit stronger behavioural responses (see D’Eath, 1998; Oliveira et al., 2000 for a critical examination of the use of videos as visual stimuli in animal behaviour research). Future studies could also combine multiple sensory modalities, e.g. by pairing a wolf image with its vocalisation and scent, to increase stimulus salience. To obtain a more comprehensive picture of how goats perceive visual stimuli, additional physiological parameters such as heart rate variability (e.g. Krause et al., 2017), salivary cortisol (e.g. Scollo et al., 2014) or eye temperature (e.g. Comin et al., 2024), could be assessed alongside looking behaviour.

## 5. Conclusion

Our study revealed that the goats’ looking behaviour was influenced by whether the presented animal (in the case of reptiles and mammals) was a two-dimensional representation of a natural predator of goats or not. Photographs of snakes attracted the goats’ visual attentionin contrast to non-predatory reptiles, and might therefore serve as potential negative stimuli in future ADAB tests. However, this increased attention towards threatening reptiles needs further validation, especially concerning the perceived emotional valence (for a critical discussion of looking time as an indicator of valence, see Crump et al., 2018). Despite the limitations of two-dimensional stimuli, they offer high experimental control, highlighting the need to find an appropriate balance between ecological validity and stimulus standardisation.

## Acknowledgements

We would like to thank the staff of the Experimental Animal Facility Pygmy Goat at the Research Institute for Farm Animal Biology in Dummerstorf, Germany, for taking care of the animals. Special thanks go to Michael Seehaus for technical support.

## Author contribution section

JD – conceptualisation, data curation, formal analysis, investigation, methodology, visualisation, writing – original draft preparation, writing – review & editing

SL – data curation, formal analysis, investigation, writing – review & editing AE – formal analysis, visualisation, writing – review & editing

CN – conceptualisation, methodology, project administration, supervision, writing – review & editing

## Data availability statement

All code and data are available online: https://doi.org/10.17605/OSF.IO/VSWU4.

## Declaration of generative AI and AI-assisted technologies in the manuscript preparation process

During the preparation of this work, the author(s) used ChatGPT/OpenAI to assist and revise R scripts. After using this tool/service, the author(s) reviewed and edited the content as needed and take full responsibility for the content of the published article.

## References

1. Allritz, M., Call, J., & Borkenau, P. (2016). How chimpanzees (*Pan troglodytes*) perform in a modified emotional Stroop task. Animal Cognition, 19(3), 435–449. 10.1007/s10071-015-0944-3

2. Alonso, M. E., González-Montaña, J. R., & Lomillos, J. M. (2020). Consumers’ Concerns and Perceptions of Farm Animal Welfare. Animals, 10(3), Article 3. 10.3390/ani10030385

3. ASAB Ethical Committee/ABS Animal Care Committee. (2024). Guidelines for the ethical treatment of nonhuman animals in behavioural research and teaching. Animal Behaviour, 207, I–XI. 10.1016/S0003-3472(23)00317-2

4. Bennett, R. (1995). The Value of Farm Animal Welfare. Journal of Agricultural Economics, 46(1), 46–60. 10.1111/j.1477-9552.1995.tb00751.x

5. Bertels, J., Bourguignon, M., De Heering, A., Chetail, F., De Tiège, X., Cleeremans, A., & Destrebecqz, A. (2020). Snakes elicit specific neural responses in the human infant brain. Scientific Reports, 10(1), 7443. 10.1038/s41598-020-63619-y

6. Bethell, E. J., Holmes, A., MacLarnon, A., & Semple, S. (2012). Evidence That Emotion Mediates Social Attention in Rhesus Macaques. PLOS ONE, 7(8), e44387. 10.1371/journal.pone.0044387

7. Brilot, B. O., & Bateson, M. (2012). Water bathing alters threat perception in starlings. Biology Letters, 8(3), 379–381. 10.1098/rsbl.2011.1200

8. Brilot, B. O., Normandale, C. L., Parkin, A., & Bateson, M. (2009). Can we use starlings’ aversion to eyespots as the basis for a novel ‘cognitive bias’ task? Applied Animal Behaviour Science, 118(3), 182–190. 10.1016/j.applanim.2009.02.015

9. Brooks, M. E., Kristensen, K., Benthem, K. J. van, Magnusson, A., Berg, C. W., Nielsen, A., Skaug, H. J., Mächler, M., & Bolker, B. M. (2017). glmmTMB Balances Speed and Flexibility Among Packages for Zero-inflated Generalized Linear Mixed Modeling. The R Journal, 9(2), 378. 10.32614/RJ-2017-066

10. Burman, O. H. P., Parker, R., Paul, E. S., & Mendl, M. (2008). A spatial judgement task to determine background emotional state in laboratory rats, *Rattus norvegicus*. Animal Behaviour, 76(3), 801–809. 10.1016/j.anbehav.2008.02.014

11. Campbell, D. L. M., Taylor, P. S., Hernandez, C. E., Stewart, M., Belson, S., & Lee, C. (2019). An attention bias test to assess anxiety states in laying hens. PeerJ, 7, e7303. 10.7717/peerj.7303

12. Chung, Y., Rabe-Hesketh, S., Dorie, V., Gelman, A., & Liu, J. (2013). A Nondegenerate Penalized Likelihood Estimator for Variance Parameters in Multilevel Models. Psychometrika, 78(4), 685–709. 10.1007/s11336-013-9328-2

13. Comin, M., Atallah, E., Chincarini, M., Mazzola, S., Canali, E., Minero, M., Cozzi, B., Rossi, E., Vignola, G., & Costa, E. D. (2024). Events with Different Emotional Valence Affect the Eye’s Lacrimal Caruncle Temperature Changes in Sheep. Animals, 14, 50. 10.3390/ani14010050

14. Crump, A., Arnott, G., & Bethell, E. (2018). Affect-Driven Attention Biases as Animal Welfare Indicators: Review and Methods. Animals, 8(8), 136. 10.3390/ani8080136

15. Cussen, V. A., & Mench, J. A. (2014). Personality predicts cognitive bias in captive psittacines, *Amazona amazonica*. Animal Behaviour, 89, 123–130. 10.1016/j.anbehav.2013.12.022

16. Da Costa, A. P., Leigh, A. E., Man, M. S., & Kendrick, K. M. (2004). Face pictures reduce behavioural, autonomic, endocrine and neural indices of stress and fear in sheep. Proceedings of the Royal Society B: Biological Sciences, 271(1552), 2077–2084. 10.1098/rspb.2004.2831

17. D’Eath, R. B. (1998). Can video images imitate real stimuli in animal behaviour experiments? Biological Reviews, 73(3), 267–292. 10.1111/j.1469-185X.1998.tb00031.x

18. Deutsch, J., Lebing, S., Eggert, A., & Nawroth, C. (2024). Goats who stare at video screens – assessing behavioural responses of goats towards images of familiar and unfamiliar con- and heterospecifics. Peer Community Journal, 4, e94. 10.24072/pcjournal.473

19. Doyle, R. E., Vidal, S., Hinch, G. N., Fisher, A. D., Boissy, A., & Lee, C. (2010). The effect of repeated testing on judgement biases in sheep. Behavioural Processes, 83(3), 349–352. 10.1016/j.beproc.2010.01.019

20. Düpjan, S., Stracke, J., Tuchscherer, A., & Puppe, B. (2017). An improved design for the spatial judgement task in domestic pigs. Applied Animal Behaviour Science, 187, 23–30. 10.1016/j.applanim.2016.11.012

21. Ede, T., & Parsons, T. D. (2023). Cognitive tasks as measures of pig welfare: A systematic review. Frontiers in Veterinary Science, 10, 1251070. 10.3389/fvets.2023.1251070

22. Evans, C. S., Evans, L., & Marler, P. (1993). On the meaning of alarm calls: Functional reference in an avian vocal system. Animal Behaviour, 46(1), 23–38. 10.1006/anbe.1993.1158

23. Finlayson, K., Lampe, J. F., Hintze, S., Würbel, H., & Melotti, L. (2016). Facial Indicators of Positive Emotions in Rats. PLOS ONE, 11(11), e0166446. 10.1371/journal.pone.0166446

24. Fraser, D. (2008). Understanding animal welfare. Acta Veterinaria Scandinavica, 50(S1), S1. 10.1186/1751-0147-50-S1-S1

25. Friard, O., & Gamba, M. (2016). BORIS: A free, versatile open-source event-logging software for video/audio coding and live observations. Methods in Ecology and Evolution, 7(11), 1325–1330. 10.1111/2041-210X.12584

26. Guimarães-Costa, R., Guimarães-Costa, M. B., Pippa-Gadioli, L., Weltson, A., Ubiali, W. A., Paschoalin-Maurin, T., Felippotti, T. T., Elias-Filho, D. H., Laure, C. J., & Coimbra, N. C. (2007). Innate defensive behaviour and panic-like reactions evoked by rodents during aggressive encounters with Brazilian constrictor snakes in a complex labyrinth: Behavioural validation of a new model to study affective and agonistic reactions in a prey versus predator paradigm. Journal of Neuroscience Methods, 165(1), 25–37. 10.1016/j.jneumeth.2007.05.023

27. Halekoh, U., & Højsgaard, S. (2014). A Kenward-Roger Approximation and Parametric Bootstrap Methods for Tests in Linear Mixed Models—The *R* Package **pbkrtest**. Journal of Statistical Software, 59(9). 10.18637/jss.v059.i09

28. Harding, E. J., Paul, E. S., & Mendl, M. (2004). Cognitive bias and affective state. Nature, 427(6972), 312–312. 10.1038/427312a

29. Harrison, R. (1964). Animal Machines: The New Factory Farming Industry. Vincent Stuart Publishers.

30. Hartig, F., Lohse, L., & leite, M. de S. (2024). DHARMa: Residual Diagnostics for Hierarchical (Multi-Level / Mixed) Regression Models (Version 0.4.7) [Computer software]. https://cran.r-project.org/web/packages/DHARMa/index.html?utm_source=chatgpt.com

31. Horstmann, G., & Herwig, A. (2016). Novelty biases attention and gaze in a surprise trial. *Attention, Perception*, & Psychophysics, 78(1), 69–77. 10.3758/s13414-015-0995-1

32. Kappel, S., Collins, S., Mendl, M. T., & Fureix, C. (2025). Looking out for danger: Theoretical and empirical issues in translating human attention bias tasks to assess animal affective states. Neuroscience & Behavioral Reviews, 169, 105980. 10.1016/j.neubiorev.2024.105980

33. Kawai, N. (2019). Do Snakes Draw Attention More Strongly than Spiders or Other Animals? In N. Kawai (Ed.), The Fear of Snakes: Evolutionary and Psychobiological Perspectives on Our Innate Fear (pp. 73–94). Springer. 10.1007/978-981-13-7530-9_5

34. Kluever, B. M., Howery, L. D., Breck, S. W., & Bergman, D. L. (2009). Predator and heterospecific stimuli alter behaviour in cattle. Behavioural Processes, 81(1), 85–91. 10.1016/j.beproc.2009.02.004

35. Krause, A., Puppe, B., & Langbein, J. (2017). Coping Style Modifies General and Affective Autonomic Reactions of Domestic Pigs in Different Behavioral Contexts. Frontiers in Behavioral Neuroscience, 11, 103. 10.3389/fnbeh.2017.00103

36. Lagisz, M., Zidar, J., Nakagawa, S., Neville, V., Sorato, E., Paul, E. S., Bateson, M., Mendl, M., & Løvlie, H. (2020). Optimism, pessimism and judgement bias in animals: A systematic review and meta-analysis. Neuroscience & Biobehavioral Reviews, 118, 3–17. 10.1016/j.neubiorev.2020.07.012

37. Langbein, J., Moreno-Zambrano, M., & Siebert, K. (2023). How do goats “read” 2D-images of familiar and unfamiliar conspecifics? Frontiers in Psychology, 14, 1089566. 10.3389/fpsyg.2023.1089566

38. Lee, C., Cafe, L. M., Robinson, S. L., Doyle, R. E., Lea, J. M., Small, A. H., & Colditz, I. G. (2018). Anxiety influences attention bias but not flight speed and crush score in beef cattle. Applied Animal Behaviour Science, 205, 210–215. 10.1016/j.applanim.2017.11.003

39. Lee, C., Verbeek, E., Doyle, R., & Bateson, M. (2016). Attention bias to threat indicates anxiety differences in sheep. Biology Letters, 12(6), 20150977. 10.1098/rsbl.2015.0977

40. Leliveld, L. M. C., Düpjan, S., Tuchscherer, A., & Puppe, B. (2016). Behavioural and physiological measures indicate subtle variations in the emotional valence of young pigs. Physiology & Behavior, 157, 116–124. 10.1016/j.physbeh.2016.02.002

41. Luikart, G., Gielly, L., Excoffier, L., Vigne, J.-D., Bouvet, J., & Taberlet, P. (2001). Multiple maternal origins and weak phylogeographic structure in domestic goats. Proceedings of the National Academy of Sciences, 98(10), 5927–5932. 10.1073/pnas.091591198

42. MacLeod, C., Mathews, A., & Tata, P. (1986). Attentional bias in emotional disorders. Journal of Abnormal Psychology, 95(1), 15–20. 10.1037//0021-843x.95.1.15

43. Mellor, D. J. (2016). Updating Animal Welfare Thinking: Moving beyond the ‘Five Freedoms’ towards ‘A Life Worth Living’. Animals, 6(3), 21. 10.3390/ani6030021

44. Mendl, M., Burman, O. H. P., Parker, R. M. A., & Paul, E. S. (2009). Cognitive bias as an indicator of animal emotion and welfare: Emerging evidence and underlying mechanisms. Applied Animal Behaviour Science, 118(3), 161–181. 10.1016/j.applanim.2009.02.023

45. Mendl, M., Burman, O. H. P., & Paul, E. S. (2010). An integrative and functional framework for the study of animal emotion and mood. Proceedings of the Royal Society B: Biological Sciences, 277(1696), 2895–2904. 10.1098/rspb.2010.0303

46. Mendl, M., Neville, V., & Paul, E. S. (2022). Bridging the Gap: Human Emotions and Animal Emotions. Affective Science, 3(4), 703–712. 10.1007/s42761-022-00125-6

47. Mendl, M., & Paul, E. S. (2020). Animal affect and decision-making. Neuroscience & Biobehavioral Reviews, 112, 144–163. 10.1016/j.neubiorev.2020.01.025

48. Miele, M., Veissier, I., Evans, A., & Botreau, R. (2011). Animal welfare: Establishing a dialogue between science and society. Animal Welfare, 20(1), 103–117. 10.1017/S0962728600002475

49. Monk, J. E., Belson, S., Colditz, I. G., & Lee, C. (2018a). Attention Bias Test Differentiates Anxiety and Depression in Sheep. Frontiers in Behavioral Neuroscience, 12(246). https://www.frontiersin.org/articles/10.3389/fnbeh.2018.00246

50. Monk, J. E., Belson, S., & Lee, C. (2019b). Pharmacologically-induced stress has minimal impact on judgement and attention biases in sheep. Scientific Reports, 9(1), Article 1. 10.1038/s41598-019-47691-7

51. Monk, J. E., Doyle, R. E., Colditz, I. G., Belson, S., Cronin, G. M., & Lee, C. (2018b). Towards a more practical attention bias test to assess affective state in sheep. PLOS ONE, 13(1), e0190404. 10.1371/journal.pone.0190404

52. Monk, J. E., Lee, C., Belson, S., Colditz, I. G., & Campbell, D. L. M. (2019a). The influence of pharmacologically-induced affective states on attention bias in sheep. PeerJ, 7, e7033. 10.7717/peerj.7033

53. Neave, H. W., Daros, R. R., Costa, J. H. C., Keyserlingk, M. A. G. von, & Weary, D. M. (2013). Pain and Pessimism: Dairy Calves Exhibit Negative Judgement Bias following Hot-Iron Disbudding. PLOS ONE, 8(12), e80556. 10.1371/journal.pone.0080556

54. Oliveira, R. F., Rosenthal, G. G., Schlupp, I., McGregor, P. K., Cuthill, I. C., Endler, J. A., Fleishman, L. J., Zeil, J., Barata, E., Burford, F., Gonçalves, D., Haley, M., Jakobsson, S., Jennions, M. D., Körner, K. E., Lindström, L., Peake, T., Pilastro, A., Pope, D. S., … Waas, J. R. (2000). Considerations on the use of video playbacks as visual stimuli: The Lisbon workshop consensus. Acta Ethologica, 3(1), 61–65. 10.1007/s102110000019

55. Olsson, K., & Hydbring-Sandberg, E. (2011). Exposure to a dog elicits different cardiovascular and behavioral effects in pregnant and lactating goats. Acta Veterinaria Scandinavica, 53(1), 60. 10.1186/1751-0147-53-60

56. Paul, E. S., Browne, W., Mendl, M. T., Caplen, G., Trevarthen, A., Held, S., & Nicol, C. J. (2022). Assessing animal welfare: A triangulation of preference, judgement bias and other candidate welfare indicators. Animal Behaviour, 186, 151–177. 10.1016/j.anbehav.2022.02.003

57. Paul, E. S., Harding, E. J., & Mendl, M. (2005). Measuring emotional processes in animals: The utility of a cognitive approach. Neuroscience & Biobehavioral Reviews, 29(3), 469–491. 10.1016/j.neubiorev.2005.01.002

58. R Core Team. (2024). R: A Language and Environment for Statistical Computing [Computer software]. R Foundation for Statistical Computing. https://www.R-project.org/

59. Raoult, C. M. C., & Gygax, L. (2018). Valence and Intensity of Video Stimuli of Dogs and Conspecifics in Sheep: Approach-Avoidance, Operant Response, and Attention. Animals, 8(7), 121. 10.3390/ani8070121

60. Rault, J.-L., Bateson, M., Boissy, A., Forkman, B., Grinde, B., Gygax, L., Harfeld, J. L., Hintze, S., Keeling, L. J., Kostal, L., Lawrence, A. B., Mendl, M. T., Miele, M., Newberry, R. C., Sandøe, P., Špinka, M., Taylor, A. H., Webb, L. E., Whalin, L., & Jensen, M. B. (2025). A consensus on the definition of positive animal welfare. Biology Letters, 21(1), 20240382. 10.1098/rsbl.2024.0382

61. Russell, J. A. (2003). Core affect and the psychological construction of emotion. Psychological Review, 110(1), 145–172. 10.1037/0033-295X.110.1.145

62. Scollo, A., Gottardo, F., Contiero, B., & Edwards, S. A. (2014). Does stocking density modify affective state in pigs as assessed by cognitive bias, behavioural and physiological parameters? Applied Animal Behaviour Science, 153, 26–35. 10.1016/j.applanim.2014.01.006

63. Shibasaki, M., & Kawai, N. (2009). Rapid detection of snakes by Japanese monkeys (*Macaca fuscata*): An evolutionarily predisposed visual system. Journal of Comparative Psychology, 123(2), 131–135. 10.1037/a0015095

64. Vögeli, S., Wolf, M., Wechsler, B., & Gygax, L. (2015). Housing conditions influence cortical and behavioural reactions of sheep in response to videos showing social interactions of different valence. Behavioural Brain Research, 284, 69–76. 10.1016/j.bbr.2015.02.007

65. Watanabe, S., Scheich, H., Braun, K., & Shinozuka, K. (2022). Visual snake aversion in Octodon degus and C57BL/6 mice. Animal Cognition, 25(1), 33–41. 10.1007/s10071-021-01527-y

