## Supplementary Material for "Goats who stare at wolves - identifying natural response stimuli for an affect-driven attention bias test in small ruminants"

**Table S1** containing date of birth, age at start of data collection and experimental pre-experience of the 30 test subjects

| Animal ID | Date of birth (dd/mm/yy) | Age at start of data collection [days] | Experimental pre-experience |
|---|---|---|---|
| A1 | 30.01.23 | 473 | Automated learning device[1] |
| A2 | 02.02.23 | 471 | Automated learning device[1] |
| A3 | 04.02.23 | 469 | Automated learning device[1] |
| A4 | 04.02.23 | 469 | Automated learning device[1] |
| A5 | 04.02.23 | 469 | Automated learning device[1] |
| A6 | 04.02.23 | 469 | Automated learning device[1] |
| B1 | 30.01.23 | 473 | Automated learning device[1] |
| B2 | 01.02.23 | 472 | Automated learning device[1] |
| B3 | 02.02.23 | 471 | Automated learning device[1] |
| B4 | 04.02.23 | 469 | Automated learning device[1] |
| B5 | 13.02.23 | 460 | Automated learning device[1] |
| B6 | 13.02.23 | 460 | Automated learning device[1] |
| C1 | 06.02.22 | 869 | Automated learning device[1]; Prosocial behaviour study[2]; Looking time paradigm[3] |
| C2 | 04.02.22 | 871 | Automated learning device[1]; Prosocial behaviour study[2]; Looking time paradigm[3] |
| C3 | 02.02.22 | 873 | Automated learning device[1]; Prosocial behaviour study[2]; Looking time paradigm[3] |
| C4 | 02.02.22 | 873 | Automated learning device[1]; Prosocial behaviour study[2]; Looking time paradigm[3] |
| C5 | 02.02.22 | 873 | Automated learning device[1]; Prosocial behaviour study[2]; Looking time paradigm[3] |
| C6 | 02.02.22 | 873 | Automated learning device[1]; Prosocial behaviour study[2]; Looking time paradigm[3] |

| | | | |
|---|---|---|---|
| D1 | 04.02.22 | 871 | Automated learning device[1]; Prosocial behaviour study[2]; Looking time paradigm[3] |
| D2 | 03.02.22 | 872 | Automated learning device[1]; Prosocial behaviour study[2]; Looking time paradigm[3] |
| D3 | 03.02.22 | 872 | Automated learning device[1]; Prosocial behaviour study[2]; Looking time paradigm[3] |
| D4 | 03.02.22 | 872 | Automated learning device[1]; Prosocial behaviour study[2]; Looking time paradigm[3] |
| D5 | 27.01.22 | 879 | Automated learning device[1]; Prosocial behaviour study[2]; Looking time paradigm[3] |
| D6 | 27.01.22 | 879 | Automated learning device[1]; Prosocial behaviour study[2]; Looking time paradigm[3] |
| E1 | 11.07.20 | 1,472 | Automated learning device[1]; Prosocial behaviour study[2]; Looking time paradigm[3] |
| E2 | 13.07.20 | 1,470 | Automated learning device[1]; Prosocial behaviour study[2]; Looking time paradigm[3] |
| E3 | 10.02.21 | 1,290 | Automated learning device[1]; Looking time paradigm[3] |
| E4 | 14.02.21 | 1,286 | Automated learning device[1]; Looking time paradigm[3] |
| E5 | 12.02.21 | 1,288 | Automated learning device[1]; Looking time paradigm[3] |
| E6 | 12.02.21 | 1,288 | Automated learning device[1]; Looking time paradigm[3] |

[1]Langbein, J., Moreno-Zambrano, M., Siebert, K. (2023). How do goats “read” 2D-images of familiar and unfamiliar conspecifics?. *Frontiers in Psychology*, 14:1089566

[2]Pahl, A., Rault, J-L., McGetrick, J., Eggert, A., Nawroth, C., Langbein, J. (2025). Do goats exhibit prosocial motivation? Insights from a novel food-giving paradigm. *Royal Society Open Science*, 12:250556

[3]Deutsch, J., Lebing, S., Eggert, A., Nawroth, C. (2024). Goats who stare at video screens – assessing behavioural responses of goats towards images of familiar and unfamiliar con- and heterospecifics. *Peer Community Journal*, 4: e94

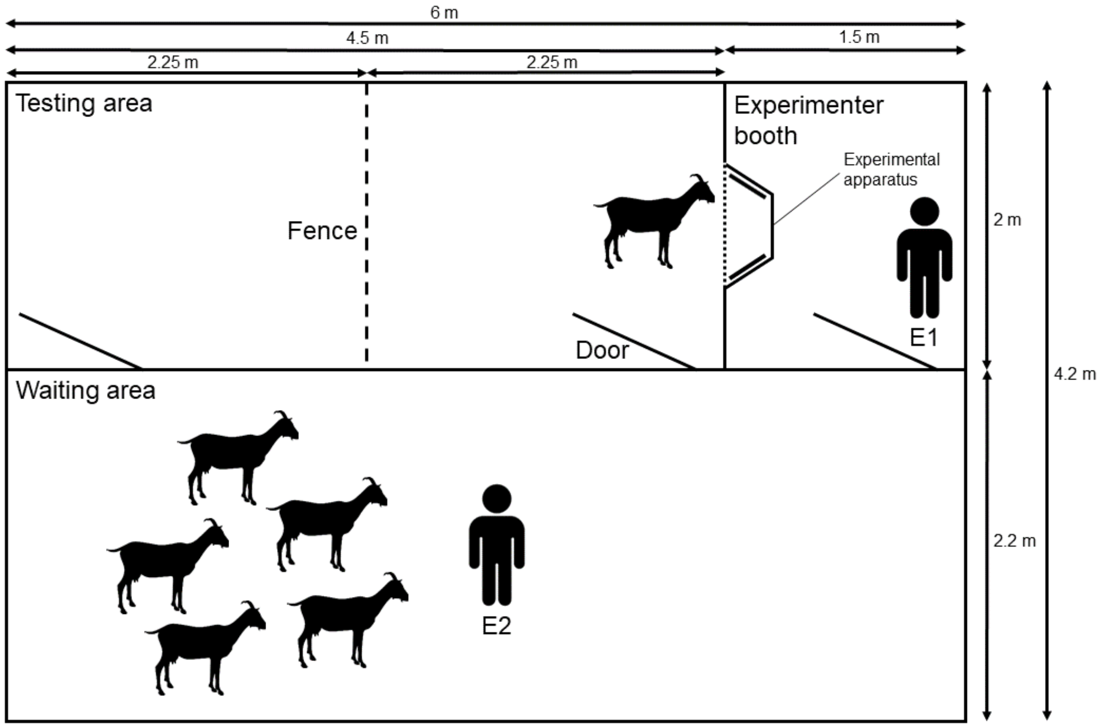


**Figure S2** Scheme of the experimental arena, including the testing area, the experimenter booth, the waiting area and the experimental apparatus

**Table S3** containing the degree of pre-experience with the looking time paradigm and the durations of the single habituation steps in the current study for each group of subjects

| **Group** | **Pre-experience with looking time paradigm** | **Duration habituation to human handlers, whole group [days]** | **Duration habituation to experimental arena, whole group [days]** | **Duration habituation to experimental apparatus, pairwise [days]** | **Duration habituation to experimental apparatus, individually [days]** |
|---|---|---|---|---|---|
| A | no | 5 | 12 | 15 | 9 |
| B | no | 5 | 12 | 15 | 9 |
| C | yes (1 previous experiment) | - | 1 | 5 | 5 |
| D | yes (1 previous experiment) | - | 1 | 5 | 5 |
| E | yes (2 previous experiments) | - | - | 3 | 5 |

**Table S4** containing the locations and the objectives of the single habituation steps

| Habituation step | Location | Objectives |
|---|---|---|
| Habituation to human handlers, whole group | Home pen | Subjects remain calm when E1 and E2 enter the home pen and can be hand-fed |
| Habituation to experimental arena, whole group | Waiting area and testing area | Subjects remain calm in experimental arena and feed out of food bowl in experimental apparatus |
| Habituation to experimental apparatus, pairwise | Testing area | Subjects remain calm in pair setting and feed out of food bowl in experimental apparatus |
| Habituation to experimental apparatus, individually | Testing area | Subjects remain calm in testing area when being alone and feed out of food bowl in experimental apparatus |

**Table S5** containing stimuli used for this study

| Photograph | Stimulus | Taxon | Predator | Image Source |
|---|---|---|---|---|
| 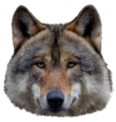 | Wolf face | Mammal | Yes | https://pixabay.com/de/photos/grauer-wolf-wolf-timberwolf-tier-7813349/ |
| 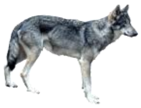 | Wolf full body | Mammal | Yes | https://pixabay.com/de/photos/wolf-nahaufnahme-natur-raubzier-3603069/ |
| 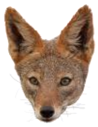 | Jackal face | Mammal | Yes | https://pixabay.com/de/photos/schakal-hund-hund-wildes-tier-8161553/ |
| 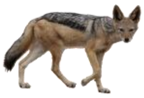 | Jackal full body | Mammal | Yes | https://pixabay.com/de/photos/silber-un-terst%C3%BCtzt-schakal-lang-4455468/ |
| 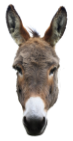 | Donkey face | Mammal | No | https://pixabay.com/de/photos/esel-lel-kendorf-grauesel-eselkopf-2095513/ |
| 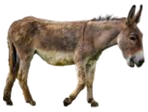 | Donkey full body | Mammal | No | https://pixabay.com/de/photos/esel-cotentin-esel-pferde-weiblich-8104157 |
| 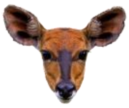 | Antelope face | Mammal | No | https://pixabay.com/de/photos/antilope-spezies-s%C3%A4ugetier-tier-7483440/ |
| 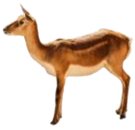 | Antelope full body | Mammal | No | https://pixabay.com/de/photos/gazelle-impala-mutter-antilope-204705/ |

| | | | | |
|---|---|---|---|---|
| 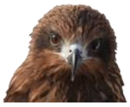 | Red kite face | Bird | Yes | https://pixabay.com/de/photos/tier-vogel-wildvogel-raptor-video-3925264/ |
| 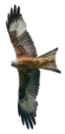 | Red kite full body | Bird | Yes | https://pixabay.com/de/photos/rotmilan-raubvogel-fliegen-2449284/ |
| 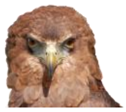 | Eagle face | Bird | Yes | https://pixabay.com/de/photos/adler-vo-gel-kopf-adlerkopf-tier-377202/ |
| 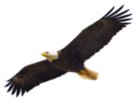 | Eagle full body | Bird | Yes | https://pixabay.com/de/photos/adler-fliege-vogel-symbol-864725/ |
| 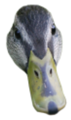 | Duck face | Bird | No | https://pixabay.com/de/photos/ente-brown-gr%C3%BCn-natur-vogel-2088263/ |
| 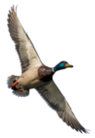 | Duck full body | Bird | No | https://pixabay.com/de/photos/ente-flug-vogel-fl%C3%BCgel-wildv%C3%B6gel-4058737/ |
| 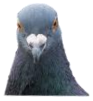 | Pigeon face | Bird | No | https://pixabay.com/de/photos/taube-vo-gel-tier-tierwelt-7230674/ |
| 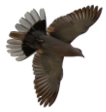 | Pigeon full body | Bird | No | https://pixabay.com/de/photos/taube-flie-gen-taube-holztaube-flug-5167481/ |
| 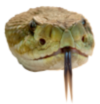 | Viper face | Reptile | Yes | https://pixabay.com/de/photos/gespren-kelte-klapperschlange-schlange-653642/ |
| 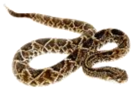 | Viper full body | Reptile | Yes | https://pixabay.com/de/photos/schlange-terrarium-bastarde-tiere-1519994/ |
| 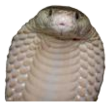 | Cobra face | Reptile | Yes | https://pixabay.com/de/photos/schlange-kobra-tier-5416747/ |
| 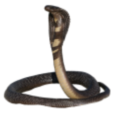 | Cobra full body | Reptile | Yes | https://pixabay.com/de/photos/schlange-kobra-klapperschlange-6285184/ |
| 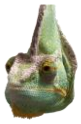 | Chameleon face | Reptile | No | https://pixabay.com/de/photos/cham%C3%A4leon-insekt-eidechse-drau%C3%9Fen-1836266/ |

| | | | | |
|---|---|---|---|---|
| 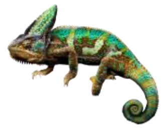 | Chameleon full body | Reptile | No | https://cdn.pixabay.com/photo/2022/03/21/14/23/chameleon-7083317_1280 |
| 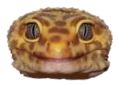 | Gecko face | Reptile | No | https://pixabay.com/de/photos/leopard-gecko-gecko-reptil-haustier-8375500/ |
| 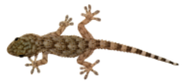 | Gecko full body | Reptile | No | https://pixabay.com/de/photos/mauergecko-tarentola-mauritanica-2656812/ |

**Table S6** containing the stimulus arrangement in the presented stimulus sets, predator stimuli in red

| Set number | Body part | Trial | Stimulus left | Stimulus right | Predator |
|---|---|---|---|---|---|
| 1 | Face | 1 | | Gecko | No |
| | | 2 | Eagle | | Yes |
| | | 3 | Antelope | | No |
| | | 4 | | Wolf | Yes |
| | | 5 | | Pigeon | No |
| | | 6 | Cobra | | Yes |
| 2 | Full body | 1 | Pigeon | | No |
| | | 2 | Wolf | | Yes |
| | | 3 | | Antelope | No |
| | | 4 | | Cobra | Yes |
| | | 5 | Gecko | | No |
| | | 6 | | Eagle | Yes |
| 3 | Face | 1 | Donkey | | No |
| | | 2 | | Viper | Yes |
| | | 3 | Chameleon | | No |
| | | 4 | | Jackal | Yes |
| | | 5 | | Duck | No |
| | | 6 | Red kite | | Yes |
| 4 | Full body | 1 | Viper | | Yes |
| | | 2 | | Duck | No |
| | | 3 | Red kite | | Yes |
| | | 4 | | Chameleon | No |

| | | | | | |
|---|---|---|---|---|---|
| | | 5 | | Jackal | Yes |
| | | 6 | Donkey | | No |
| 5 | Face | 1 | Gecko | | No |
| | | 2 | | Eagle | Yes |
| | | 3 | | Antelope | No |
| | | 4 | Wolf | | Yes |
| | | 5 | Pigeon | | No |
| | | 6 | | Cobra | Yes |
| 6 | Full body | 1 | | Pigeon | No |
| | | 2 | | Wolf | Yes |
| | | 3 | Antelope | | No |
| | | 4 | Cobra | | Yes |
| | | 5 | | Gecko | No |
| | | 6 | Eagle | | Yes |
| 7 | Face | 1 | | Donkey | No |
| | | 2 | Viper | | Yes |
| | | 3 | | Chameleon | No |
| | | 4 | Jackal | | Yes |
| | | 5 | Duck | | No |
| | | 6 | | Red kite | Yes |
| 8 | Full body | 1 | | Viper | Yes |
| | | 2 | Duck | | No |
| | | 3 | | Red kite | Yes |
| | | 4 | Chameleon | | No |
| | | 5 | Jackal | | Yes |
| | | 6 | | Duck | No |

**Table S7** containing subject stimulus set assignment, subject A2 needed to be excluded after session 2

| Subject | Session 1 | Session 2 | Session 3 | Session 4 |
|---|---|---|---|---|
| A1 | Set 1 | Set 2 | Set 3 | Set 4 |
| A2 | Set 4 | Set 3 | - | - |
| A3 | Set 3 | Set 4 | Set 1 | Set 6 |
| A4 | Set 2 | Set 5 | Set 8 | Set 7 |
| A5 | Set 5 | Set 6 | Set 7 | Set 8 |
| A6 | Set 8 | Set 7 | Set 6 | Set 1 |
| B1 | Set 8 | Set 7 | Set 2 | Set 5 |
| B2 | Set 7 | Set 8 | Set 5 | Set 6 |
| B3 | Set 6 | Set 1 | Set 8 | Set 7 |
| B4 | Set 3 | Set 4 | Set 1 | Set 2 |
| B5 | Set 2 | Set 5 | Set 4 | Set 3 |
| B6 | Set 1 | Set 6 | Set 3 | Set 4 |
| C1 | Set 3 | Set 4 | Set 1 | Set 2 |
| C2 | Set 2 | Set 5 | Set 4 | Set 3 |
| C3 | Set 1 | Set 6 | Set 3 | Set 4 |
| C4 | Set 8 | Set 7 | Set 2 | Set 5 |
| C5 | Set 7 | Set 8 | Set 5 | Set 6 |
| C6 | Set 6 | Set 1 | Set 8 | Set 7 |
| D1 | Set 2 | Set 5 | Set 8 | Set 7 |
| D2 | Set 5 | Set 6 | Set 7 | Set 8 |
| D3 | Set 8 | Set 7 | Set 6 | Set 1 |
| D4 | Set 1 | Set 2 | Set 3 | Set 4 |
| D5 | Set 4 | Set 3 | Set 2 | Set 5 |
| D6 | Set 3 | Set 4 | Set 1 | Set 6 |
| E1 | Set 4 | Set 3 | Set 2 | Set 5 |
| E2 | Set 3 | Set 4 | Set 1 | Set 6 |
| E3 | Set 2 | Set 5 | Set 8 | Set 7 |
| E4 | Set 5 | Set 6 | Set 7 | Set 8 |
| E5 | Set 8 | Set 7 | Set 6 | Set 1 |
| E6 | Set 1 | Set 2 | Set 3 | Set 4 |

**Table S8** containing the model estimates, lower confidence intervals and upper confidence intervals for each presented stimulus class of the GLMM for RLD_Out

| Category | Taxon | Body part | Estimate | Lower CI | Upper CI |
|---|---|---|---|---|---|
| Non-predator | Bird | Face | 0.362 | 0.277 | 0.456 |
| Predator | Bird | Face | 0.36 | 0.277 | 0.453 |
| Non-predator | Bird | Body | 0.258 | 0.184 | 0.349 |
| Predator | Bird | Body | 0.296 | 0.212 | 0.395 |
| Non-predator | Mammal | Face | 0.385 | 0.298 | 0.48 |
| Predator | Mammal | Face | 0.385 | 0.294 | 0.484 |
| Non-predator | Mammal | Body | 0.319 | 0.236 | 0.416 |
| Predator | Mammal | Body | 0.362 | 0.275 | 0.459 |
| Non-predator | Reptile | Face | 0.313 | 0.237 | 0.401 |
| Predator | Reptile | Face | 0.337 | 0.253 | 0.432 |
| Non-predator | Reptile | Body | 0.289 | 0.21 | 0.383 |
| Predator | Reptile | Body | 0.355 | 0.271 | 0.449 |
